# Computationally guided insights into substrate recognition and specificity of Alg1

**DOI:** 10.1101/2023.12.27.573458

**Authors:** Cheng Yu, Fangyuan Long, Ruijie Li, Yi Ding

## Abstract

The assembly of N-linked glycans for the N-glycosylation, which is an important protein modification process, starts with the consecutive synthesis of the dolichol-linked oligosaccharide (DLO) precursor in the endoplasmic reticulum (ER). Asparagine-linked glycosylation protein 1 (Alg1) is the first mannosyltransferase (MTase) in DLO synthesis, which transfers a mannose residue from the sugar donor GDP-mannose (GDP-Man) to the substrate dolichol pyrophosphate-GlcNAc2 (Dol-PP-GlcNAc2) to form a β-1,4 glycosidic bond. However, little is known about its process of substrate recognition and the mechanism for its substrate specificity. Here, the models of large-scale systems in solution and membrane environments were built, revealing the substrate recognition and specificity mechanism of Alg1 based on extensive all-atom microsecond molecular dynamics (MD) simulations. In this study, the unique non-natural donor recognition of *Sc*Alg1 and its mechanisms was reported. Further, the cooperation of the hydrophobic region and lysine in substrate recognition was revealed for both *Sc*Alg1 and *Hs*Alg1. Moreover, the functional roles of these sites were confirmed via mutant activity assays in *vitro*. Additionally, the substrate binding process of Alg1 was simulated, elucidating the effect of dolichol on substrate specificity. This work reports the substrate recognition of Alg1 and reveals mechanisms for its sugar donor and substrate selectivity, which could further aid in mechanistic studies of the DLO biosynthetic process and drug development targeting Alg1.

## 1. Introduction

In eukaryotic cells, N-glycosylation is a significant post-translational modification occurring in most proteins [1], playing a crucial role in physiological processes such as signal transduction and immune response [2]. This modification begins in ER [3] and is completed in the Golgi apparatus [4], encompassing three main subtypes: hybrid, complex, and high mannose type, all sharing a common pentasaccharide core Man3-GlcNAc2-Asn [5]. The donors for N-glycosylation, DLOs, are synthesized in *vivo* by a series of glycosyltransferases (GTs) known as Alg [6]. Key enzymes like Alg7, Alg13/14, Alg1, and Alg2, are MTases of the B-superfamily (GT-Bs), responsible for assembling Dol-PP-GlcNAc2-Man3 [5]. These GTs, using nucleotide sugars as donors, sequentially add monosaccharide residues on Dol-PP [7]. The synthesis of DLO is orchestrated by the substrate recognition dynamics of these enzymes, facilitating sequential biosynthesis based on their structure. Therefore, exploring the substrate recognition mechanisms of these GTs is essential for understanding the N-glycosylation process [8].

Alg1, the initial MTase in the synthesis pathway of DLOs, utilizes GDP-Man as the sugar donor, adding mannose onto Dol-PP-GlcNAc2 via a beta-1,4 linkage [5,9]. Belonging to the GT33 family, Alg1 is a typical GT-B protein [10]. Past research mainly focused on yeast Alg1 and used the natural substrate Dol-PP-GlcNAc2 to test the activity [11]. Later, a DLO analogue, phytanyl-pyrophosphoryl-α-N, N′-diacetylchitobioside (PPGn2), was synthesized as the substrate for *Sc*Alg1 [12], with its activity confirmed via UPLC-MS [13]. The physiological significance of *Sc*Alg1’s N-terminal alpha helix has been demonstrated to aid its ER membrane localization [14]. Mutations in *Hs*Alg1 lost parts of its activity, leading to *ALG1*-congenital disorder of glycosylation (Alg1-CDG), which was recognized as a major cause of CDG [15]. Numerous Alg1-CDG cases have been reported in studies involving *Hs*Alg1 mutants [16–18].

As a membrane protein with multiple transmembrane domains, Alg1’s substrate specificity and relatively complex structure pose challenges for research. Current research on Alg1 relied on reactivation tests with mutants in *vivo* [14] and activity tests with different substrates in *vitro*. Although the β-1,4 MTase activity of Alg1 has been quantitatively detected [13], its key recognition sites and molecular mechanisms remain unknown. Regarding protein structure, only the structures of Alg13 [19] and Alg6 [20] among these Alg proteins have been elucidated. While Alg13 will form complexes with Alg14 for full functionality [21], Alg6 belongs to the GT-C family [22]. Thus, for Alg1 and even other Alg proteins, reports on their substrate recognition and catalytic mechanisms were scarce, which also significantly hindered research on the pathogenesis of CDG diseases.

The complex natural extraction and chemical synthesis of DLO significantly hindered research into Alg1 in *vitro*. Dol-PP-GlcNAc2 is the natural substrate of Alg1, with which the dolichol tail is between fourteen to twenty-one isoprenyl units long [23]. Although PPGn2 or other synthetic LLO analogues can serve as the substrate for *Sc*Alg1, its activity was lower than that with natural substrates [24]. Due to its membrane-dependent transglycosylation activity and its high substrate preference for natural substrate, the function of *Hs*Alg1 in *vitro* has never been reported. Studying these molecular mechanisms can provide crucial guidance for donor and substrate design [25], which is vital for understanding the process of N-glycosylation, the synthesis of oligosaccharides [26]. In the field of biology, computational simulations can serve as a crucial tool, enabling an in-depth analysis and comprehension of the atomic-level structural and dynamical properties of a broad spectrum of essential biomolecules found within living [27].

In this study, we reported the substrate recognition mechanisms and substrate specificity of Alg1 from *S. cerevisiae* and *H. sapiens*. Predicted by AlphaFold2 [28], structures of Alg1 were obtained and substrate complexes were placed in *vitro* reactions [13] and physiological environments separately to simulate their substrate recognition processes. Compared to past research, this study reveals Alg1’s ability to recognize both natural and non-natural sugar donors, establishing the predicted binary complex of *Sc*Alg1 with its sugar donor, and elucidating its sugar donor promiscuity [29] at a molecular level. Furthermore, molecular simulations were employed to elucidate the key binding sites and mechanisms of substrate interaction for Alg1, conducted in two distinct environments: an aqueous solution to represent reaction conditions in *vitro* and on the ER membrane to simulate physiological contexts. Utilizing soluble *Sc*Alg1-TM as the starting point [22], in *vitro,* enzymatic assays were conducted to verify its sugar donor and substrate recognition mechanisms. Particularly, long-duration simulations of *Hs*Alg1 with natural substrate and substrate with short-chain were conducted respectively for the first time, offering a reliable molecular perspective on substrate binding and selectivity. Using the computer molecular simulation, this study provides the pathway for research on membrane-bound glycosyltransferases like Alg1 and the potential theoretical base for the treatment of CDG.

## 2. Materials and methods

### 2.1 Plasmids, strains

The primers utilized in this study were comprehensively catalogued (Supplementary Table I). The plasmids were synthesized employing standard protocols in molecular biology. Site-directed mutagenesis of various *Sc*Alg1 mutants was accomplished via fusion-PCR, with subsequent confirmation through DNA sequencing. All variants of Alg1, including its mutant forms, were engineered to be expressed under the control of the galactose operon, featuring an N-terminal 6-His tag. For heterologous expression in *Escherichia coli Rosetta* (DE3) cells, Alg1 mutants were strategically cloned into the pET28a vector.

### 2.2 Protein expression and purification

*Sc*Alg1 proteins (*Sc*Alg1-TM) were expressed and purified as described previously [14]. In this experimental protocol, *Escherichia coli Rosetta* (DE3) cells harboring the pET28a-*6His*-*ALG1*-TM plasmid were cultured in Terrific Broth (TB) medium at 37°C until the OD_600_ reached approximately 0.9. This culture was then shifted to 16°C, and protein expression was induced by adding 0.1 mM isopropyl β-D-thiogalactopyranoside (IPTG) followed by an 18-hour incubation. Cells were harvested, resuspended in Buffer I (50 mM Tris-HCl, pH 8.0, 150 mM NaCl), and lysed via ultrasonication on ice. The cell lysates were then centrifuged (4,000 × g, 30 min) to remove cellular debris. The supernatant was treated with 1% v/v Triton X-100, stirred gently at 4°C for one hour, and centrifuged again (15,000 × g, 30 min) to eliminate insoluble material. The Alg1 mutant proteins were purified from the resulting supernatant using Ni-NTA affinity chromatography (GE healthcare, Buckinghamshire, UK). SDS-PAGE was used to test their purity (Fig. S3). Post-purification, the proteins were dialyzed overnight at 4°C in dialysate buffer (25 mM Tris-HCl, pH 8.0, 100 mM NaCl) and subsequently concentrated to 1 mg/mL using an Amicon^®^ Ultra-15 centrifugal filter unit with a 10 kDa NMWL membrane (Millipore, MA, USA).

### 2.3 Enzymatic synthesis of non-natural sugar donors

GDP-Man, GDP-glucose, GDP-fucose, UDP-glucose and UDP-GlcNAc were purchased from Glycolipidpharm. The non-natural sugar donors including GDP-altrose and GDP-glucose are enzymatically synthesized as reported [30]. Briefly, sugars were added into the systems with three enzymes (Nahk, pFManc and EcpPA) and the buffer (100 mM GTP, 25 mM Tris-HCl (pH 8.0), 150 mM NaCl (Sigma-Aldrich, MO, USA)) for 72 h at 3□[ (Fig. S4, S5, S6). After the reaction, molecular sieve gel column chromatography was used to purify the sugar donors (Fig. S2).

### 2.4 Activity assay in *vitro*

An MS-based quantitative assay was used to measure the enzymatic activity of Alg1 [13]. In this assay, the reaction mixture, with a final volume of 50 μL, comprised the following components: 20 mM Tris-HCl (pH 7.2), 1 mM dithiothreitol (DTT), 150 mM EDTA, 0.13% NP-40, 50 μM PPGn2, 2 mM GDP-Man, 10 mM MgCl_2_, and 4 ng of the mutant Alg1 protein. The enzymatic reaction was conducted at 30°C for 30 minutes and subsequently terminated by heating at 100°C for 2 minutes. Post-reaction, the mixture was lyophilized and then reconstituted in 0.2 mL of 40 mM HCl [H_2_O/MeOH (4:1)], followed by incubation at 100°C for 1 hour to release glycans from pyrophosphate phytanol (PP-Phy). The aqueous fraction was desalted using solid-phase extraction with 1 mL of Supelclean^TM^ ENVI-Carb^TM^ SPE tube (Sigma, MO), pre-equilibrated with 2% acetonitrile. The column was washed with 10 mL of 2% acetonitrile, and the oligosaccharides were eluted using 3.0 mL of 25% acetonitrile, then lyophilized. The desalted oligosaccharides were analyzed using a Dionex Ultimate 3000 UPLC system (Thermo Scientific, MA), under the following conditions: Waters Acquity UPLC BEH Amide Column (1.7 μm, 2.1 × 100 mm); eluent A being CH_3_CN, and eluent B H_2_O; with a gradient of 20% B for 0–2 min, increasing to 50% B over 2–15 min, and holding at 50% B for 15–18 min at a flow rate of 0.2 mL/min. The eluate’s ESI-MS was recorded on a TSQ Quantum Ultra mass spectrometer (Thermo Scientific, MA) in the mass range of 400–800 m/z in positive mode. The oligosaccharide transfer rate was quantified by analyzing the peak intensities in the UPLC–ESI-MS data using Xcalibur software (Version 2.0, Thermo Scientific, USA).

### 2.5 Modelling and molecular docking

The structures of *Sc*Alg1 and *HsAlg1 were* predicted by AlphaFold2 [28]. The structure of *Sc*Alg1-TM was based on the structure of *Sc*Alg1.The binary and ternary complexes were built by PyMOL. All the molecular docking was carried by Maestro Glide. Due to the limitation of Maestro, complex ligands like Dol16-PP-GlcNAc2 were not used for molecular docking. The PPGn2 with C20 lipid chain was used in activity tests of *Sc*Alg1-TM and the PPGn2 with C15 lipid chain was used in molecular dockings and simulations (Fig. S1). The visualization was carried out by PyMOL and VMD.

### 2.6 Membrane system built

CHARMM-GUI was used to build the system including protein complexes and ER membrane. Briefly, the OPM server provided pre-oriented protein coordinates concerning the membrane normal [31,32]. The CHARMM-GUI membrane builder was used to insert the protein motifs into a bilayer membrane. The bilayer system was composed of 224 POPC, 88 DOPE, 44 SAPI and 28 cholesterol molecules per leaflet [33] in transferrable intermolecular potential with three points (TIP3P) water and 100 mM NaCl and 10 mM MgCl_2_.

### 2.7 Molecular dynamics (MD) simulations

The all-atom simulations were performed with GROMACS-2021.6 and the Charmm36m forcefield. All systems for *Sc*Alg1-TM were solvated in a cubic box of 103 × 103 × 103 Å^3^ with TIP3P water molecules and electroneutralized with 100 mM NaCl and 10 mM MgCl_2_ counterions. All systems for *Hs*Alg1 and full-length *Sc*Alg1 were solvated in a cubic box of 155 × 155 × 130 Å^3^ with membrane, TIP3P water molecules and electroneutralized with 100 mM NaCl and 10 mM MgCl_2_ counterions (Fig. S9). Each system contained from 100,000 to 600,000 atoms. Energy minimization was executed utilizing the steepest descent method. The system underwent initial equilibration for 1 ns under constant volume and temperature conditions (NVT ensemble), followed by a subsequent 1 ns equilibration under constant pressure and temperature conditions (NPT ensemble). Temperature and pressure regulation during these phases were achieved using the Berendsen thermostat and barostat respectively. Production molecular dynamics simulations were performed at temperatures of 303.15 K or 310.15 K and a pressure of 1 atm (NPT ensemble), spanning 100 ns to 1000 ns. These conditions were maintained using the v-rescale method for temperature and Berendsen coupling for pressure regulation. Short-range nonbonded interactions were computed with a cutoff distance of 1.0 nm.

### 2.8 Root-mean-square deviation (RMSD), Root-mean-square fluctuation (RMSF)

Ligand interaction analyses were conducted utilizing the AutodockVina and Maestro. To quantify the proximity between protein residues and ligands throughout the simulations, minimum distance calculations were performed in GROMACS. The root-mean-square deviations (RMSDs) and root-mean-square fluctuations (RMSFs) about the backbone and Cα atoms, respectively, were computed over the timeframe using GROMACS, averaging results from three independent simulations. The plots illustrating RMSD and Cα RMSF were generated using Python, facilitating visual data interpretation. Variations in RMSF (ΔRMSF) for each experimental condition were calculated relative to the mean apo Cα RMSF, with shaded regions in the plots indicating the standard deviation differences.

### 2.9 Ligand binding free energy

The Lennard-Jones potential was calculated by gmx commands. The binding energy including VDW and polar energy was implemented by the gmx_MMPBSA [34]. The fully ligated protein/substrate complex trajectory was used to determine the energetics of each step. From the substrate/product complex trajectory, 100 frames evenly spread across the entire trajectory were chosen.

## 3. Results and discussion

### 3.1 Recognition of non-natural sugar donors and production of non-natural products by *Sc*Alg1-TM

To investigate the sugar donor specificity of yeast Alg1 in *vitro*, the transmembrane domain (TMD) truncated *Sc*Alg1 (*Sc*Alg1-TM) was soluble expressed [14]. When searching for the catalytic mechanism of *Sc*Alg1, the non-natural donors were used to challenge its donor specificity (Fig. 1A). Referring to the enzymatic synthesis of GDP-mannose [30], GDP-D-altrose (GDP-Alt) and GDP-glucose (GDP-Glc) were synthesized (Fig. 1B). With these non-natural donors, the glycosyltransferase activity of *Sc*Alg1 was tested in *vitro* and the products were detected by UPLC-MS [6].

**Figure 1.**
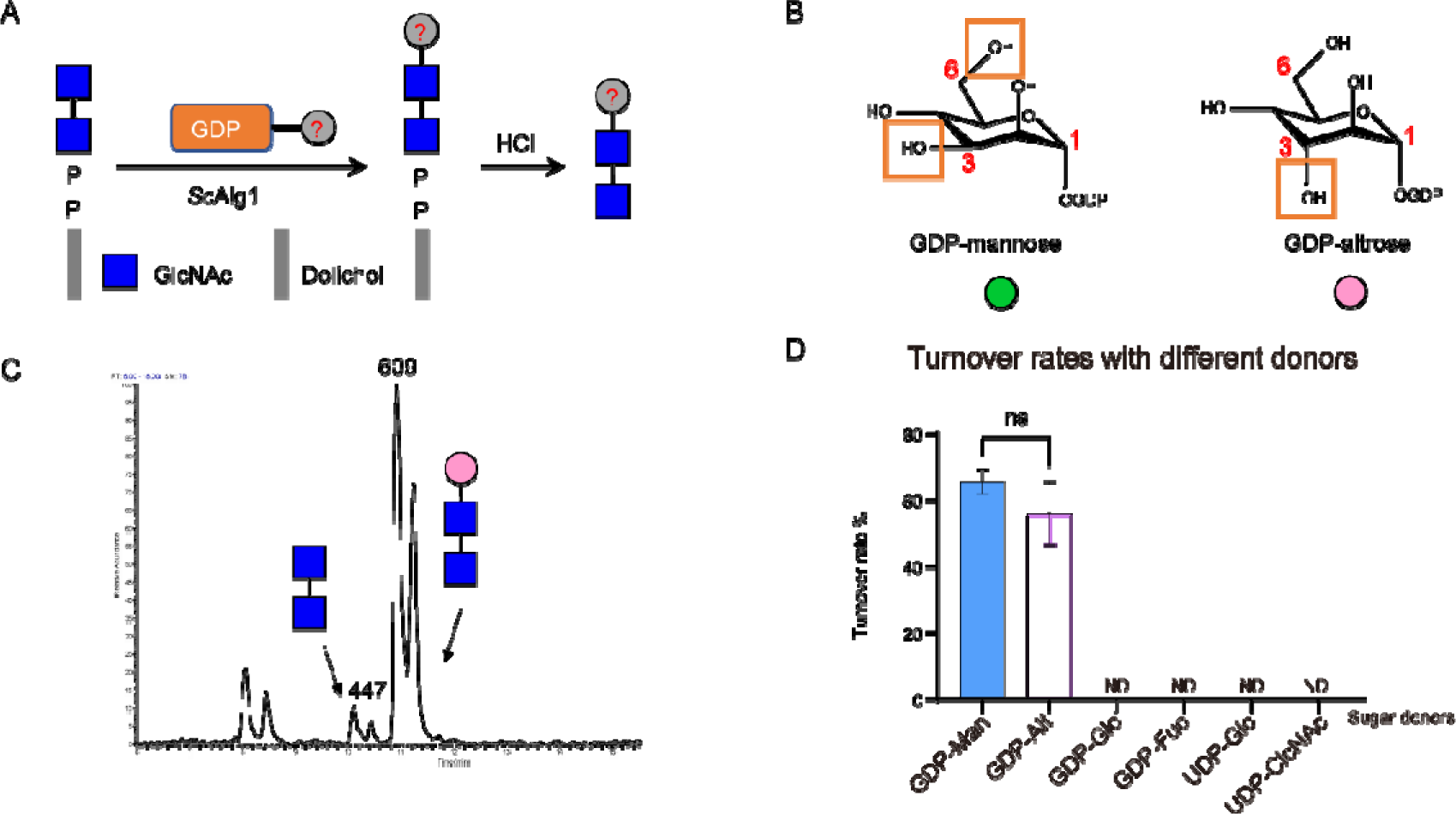
Donor specificity of *Sc*Alg1-TM. (A) Glycosyltransferase activity and the product detection of *Sc*Alg1. (B) Structure of the GDP-mannose and GDP-altrose. (C) Detection of the products of *Sc*Alg1-TM using GDP-Alt by UPLC-MS. (D) The turnover rate of *Sc*Alg1-TM in 30 minutes with different sugar donors. (ND: no turnover rate detected.)

As the MS results, *Sc*Alg1-TM could recognize these non-natural sugar donors and produce Dol-PP-GlcNAc2Alt in 2 hours (Fig. 1C) while GDP-Glc and some other sugar donors showed no activity. In the experiment in *vitro*, the unusual non-natural sugar donor recognition specificity drew attention. After strictly limiting the reaction time and conditions, GDP-Alt showed a similar turnover rate with GDP-Man (Fig. 1D). In terms of molecular structure, altrose is a C3 isomer of mannose (Fig. 1B, S2), and it was speculated that the difference in the donor preferences may come from the enzyme’s recognition of the donor hydroxyl group [35].

Based on the aforementioned analyses, it was found that *Sc*Alg1 could use non-natural sugar donors and produce non-natural substrates for the following glycosyltransferase. While substrate and donor promiscuities of GTs were observed to produce different small molecule natural products [36], it was the first time that the non-natural sugar was introduced into glycosylation of protein by enzyme reaction, even with a similar preference for natural donor. Thus, with the completely different structure of the sugar branch, the characteristics of enzymes with N-glycosylation may change greatly in *vivo*.

### 3.2 Molecular mechanism of donor recognition of *Sc*Alg1-TM

To figure out the molecular mechanism of donor recognition, molecular docking and dynamic simulation was carried out. The model of *Sc*Alg1 was predicted (Fig. 2A) by AlphaFold2 [28]. The TMD in the N-terminal was truncated to make it soluble expressed as the enzyme activity test in *vitro* [14]. Subsequently, molecular docking was performed to construct the binary complex of *Sc*Alg1 and its natural sugar donor, which is GDP-mannose (Fig. 2B). Then the system composed of the protein-donor complex and the 100 mM NaCl_2_ solution was built. The simulation of *Sc*Alg1 and its donor was carried out without limitation and the typical frames were extracted and parsed [37].

**Figure 2.**
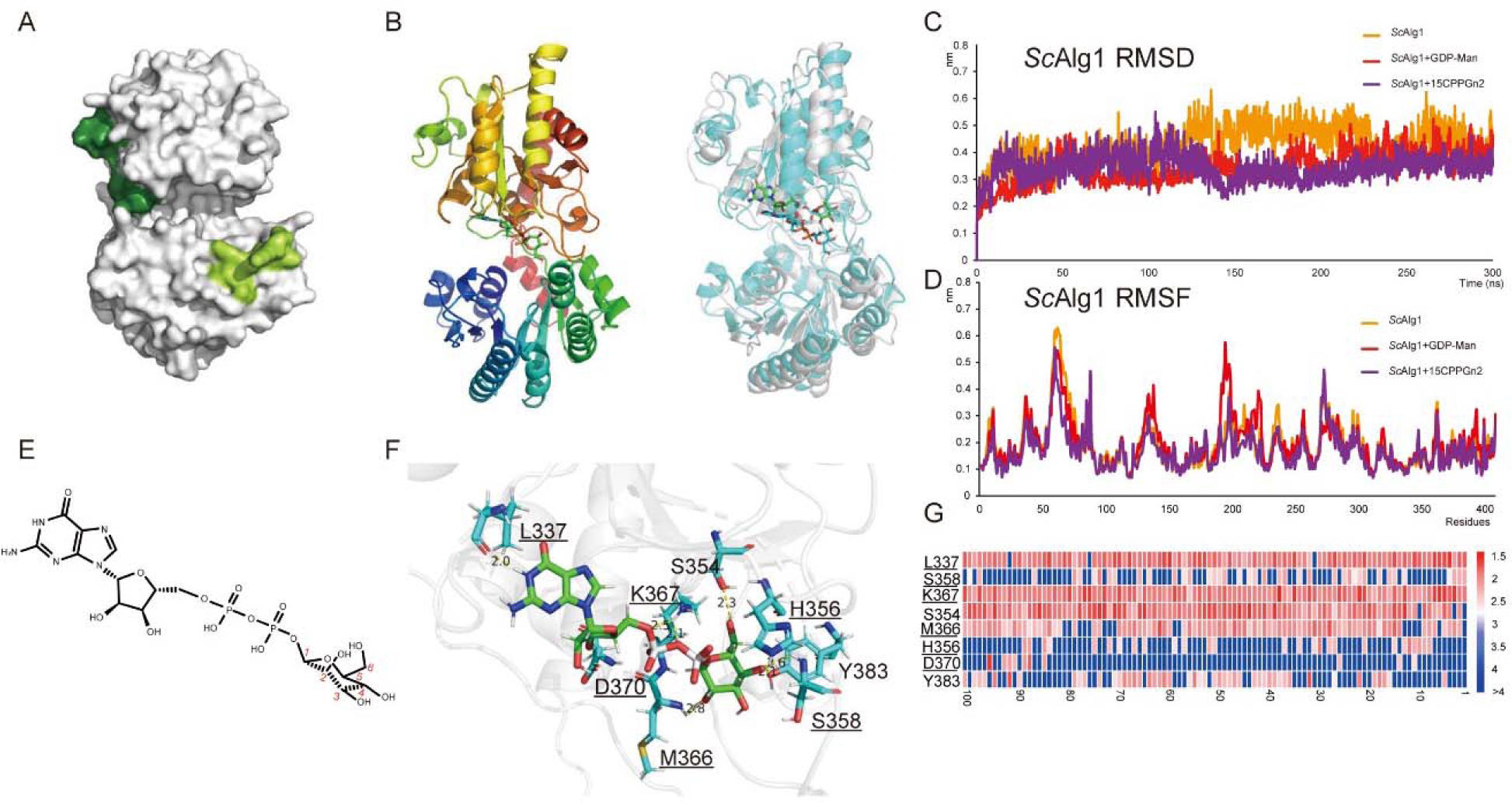
The simulation of *Sc*Alg1-TM shows the key interactions for donor recognition. (A) The predicted model of *Sc*Alg1-TM with inaccurate parts is marked green. (B) The structure of the complex *Sc*Alg1-TM/GDP-Man and the structure before (blue) and after (grey) MD. (C) The RMSD of the molecular dynamic simulation. (D) The RMSF of the molecular dynamic simulation. (E) The structure of GDP-Man. (F) The key sites of the interactions between *Sc*Alg1-TM and GDP-Man after MD. (F) The statistics of the distances between residues and GDP-Man.

The position of the sugar donor remained stable during the 300 ns MD simulation (Fig. 2C, 2D). Upon analyzing the interactions between the substrate and the protein, it was observed that several amino acids in the C-terminal region of *Sc*Alg1 interacted with GDP-Man, particularly with the sugar moiety (Fig. 2E). The donor recognition region was in the C-terminal side of the pocket of the two Rossmann folds (Fig. 2F, 2G). The pyrophosphate moiety of GDP-Man engaged in a salt-bridge interaction with the lysine at position 367 (K367), while the mannose moiety’s hydroxyl groups formed polar interactions with surrounding amino acid residues. These interactions included methionine at position 366 (M366) with the C2 hydroxyl group (C2-OH), a cluster of histidine at position 356 (H356), serine at position 358 (S358), and tyrosine at position 383 (Y383) with the C4-OH, and serine at position 354 (S354) with the C6-OH. The polar interactions facilitated the identification of sugar donors [38].

The C3-OH of the sugar was free, which explained the phenomenon that GDP-Alt showed similar activity when it replaced the natural donor GDP-Man. Also, the C6-OH of the mannose interacted with the side chain of S354, which is the non-conversed site. The combination of these polar interactions between the donor and *Sc*Alg1-TM showed a relatively strict donor recognition mechanism.

The docking and simulation helped find the key sites for donor recognition, which revealed that the absence of the interaction with the C3-hydroxyl group led to this phenomenon. Thus, it was speculated that the C3 group on the sugar scaffold was relatively flexible and it will come to be a potential site if special sugar donors targeting Alg1 need to be designed.

### 3.3 The hydrophobic region promotes the substrate recognition of *Sc*Alg1

The catalytic mechanisms of GT-B enzymes were intricate, initiating the binding of the sugar donor to the substrate in the initial step [39]. To analyze the molecular mechanism of substrate recognition of *Sc*Alg1-TM, the complex of enzyme and PPGn2 was built by molecular docking [40]. Then the complex is solvated for molecular dynamics simulation.

The N-terminal domain was composed of several α-helixes and the three helixes form a hydrophobic region [41] which contains a hydrophobic channel (Fig. 3A, 3B). As analyzed before, the third α-helix in the N-terminal is amphipathic but is not a transmembrane domain [14]. After molecular docking, it was found that the three helixes (the 3^rd^, 4^th^ and 5^th^ helix) formed a hydrophobic region and the lipid chain was inside the groove (Fig. 3C, 3D). During the all-atoms simulation, the substrates were kept in the surroundings of this region with the sugar parts oscillating between the catalytic pocket and the solution.

**Figure 3.**
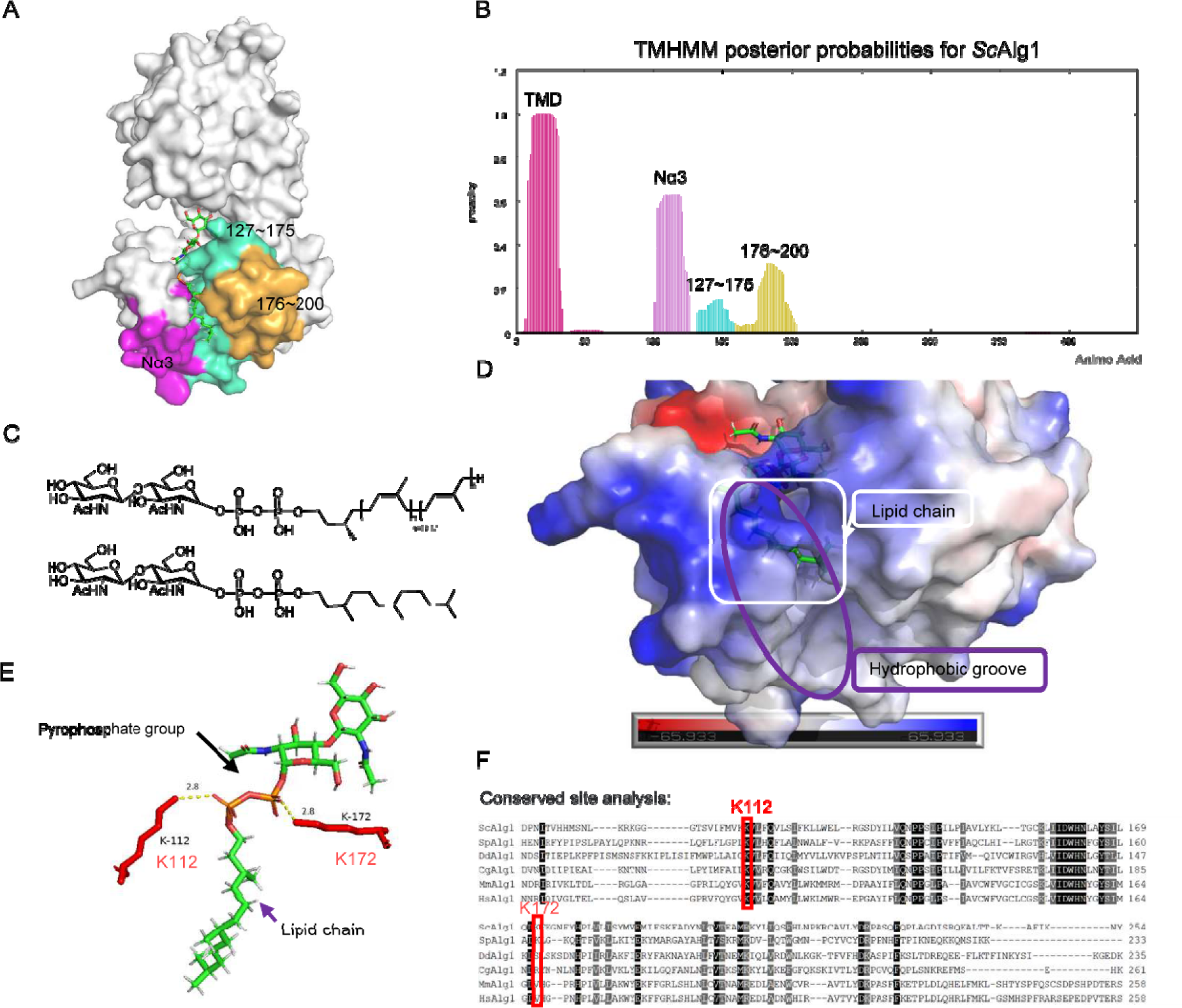
The key region and sites for the substrate recognition of *Sc*Alg1-TM. (A) The special substrate recognition region of *Sc*Alg1-TM is composed of these three α-helixes which are highlighted with distinct colours: light magenta, cyan, and sand for visualization. (B) The hydrophobicity prediction of *Sc*Alg1 by TMHMM. (C) The natural substrate (above) and the alternative substrate (below). (D) The region for the substrate recognition and the location of the lipid chain. (E) The salt bridges between the lysines and the substrate. (F) The conserved site analysis of the Alg1 from different species and conserved residue K112.

After analyzing the distance between residues and substrate molecular during simulation (Fig. S7), the lysine residue located at the N-terminal end of the 3rd helix, which was also located near the catalytic pocket, showed an irreplaceable role in substrate recognition as expected [42]. In the case of molecular docking, K112 and K172 formed salt bridges together with the phosphodiesters (Fig. 3E) and provided the energy to attract the substrate to get it into the right position [43], while the K112 is conversed and K172 is non-conversed (Fig. 3F).

In the simulation, the conversed lysine showed eternal interaction with the substrate while K172 showed low interactions with the substrate. To figure out the role of the two lysine residues in substrate recognition, enzyme activity was performed in *vitro*.

### 3.4 Enzyme activity of key sites mutants in *vitro*

To verify the key sites for donor and substrate recognitions, several predicted key sites were mutated to their allelic sites for activity testing in *vitro* (Fig. 3F, 4A). For donor recognition, K367I showed no activity, which means the residues are irreplaceable. H356 was predicted to be related to the complexation of metal ions, and the mutant H356A lost its activity, just as the test [13] without adding Mg^2+^(Fig. 4B).

**Figure 4.**
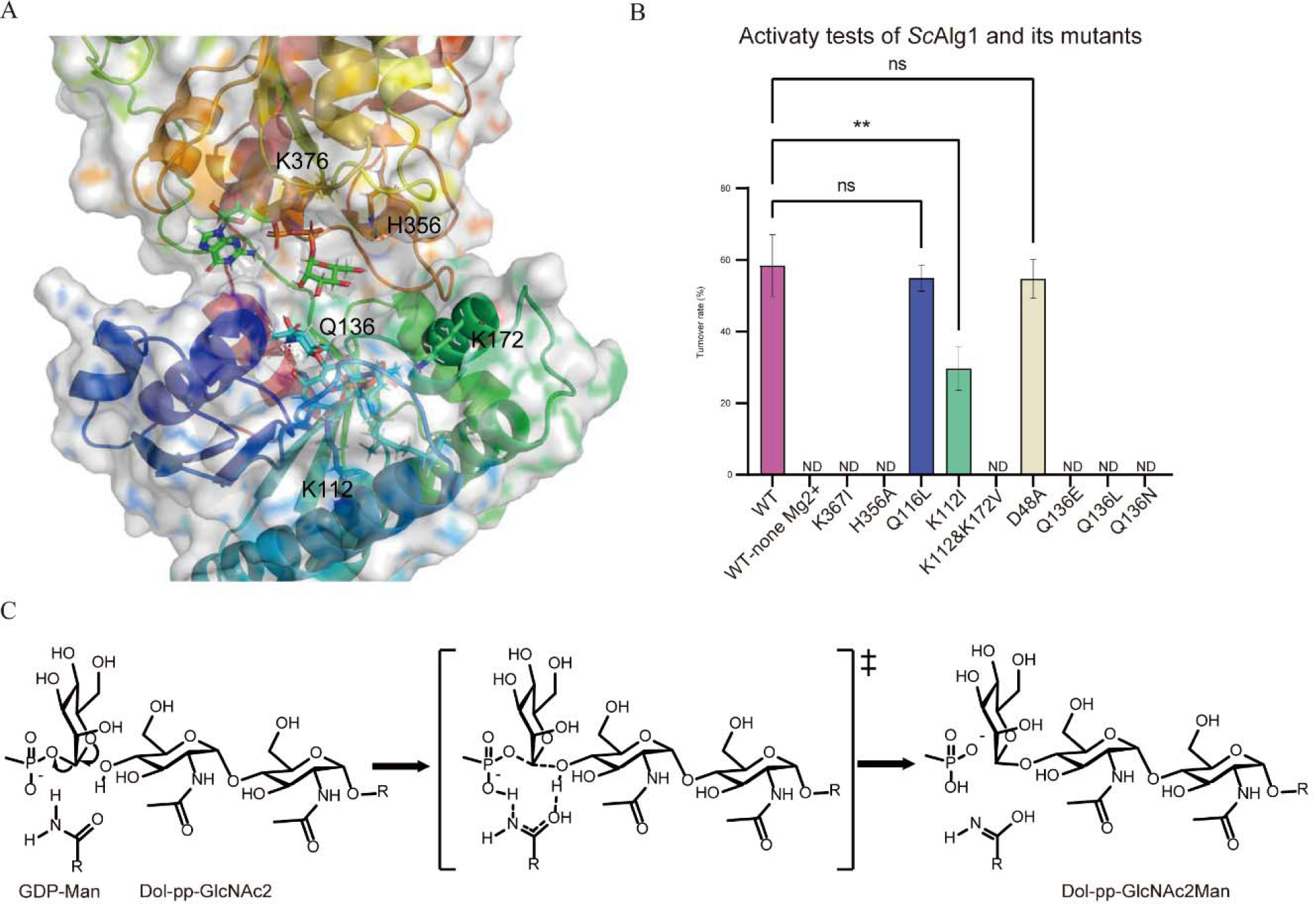
The key sites for substrates recognition and catalytic mechanism of *Sc*Alg1. (A)The key sites of *Sc*Alg1 substrates recognition. (B) The turnover rates of different mutants. (C) The predicted catalytic mechanism of *Sc*Alg1.

K112 was mutated to isoleucine which is similar to lysine, with K172 was mutated to valine which is the allelic site of *Hs*Alg1. In the enzyme activity test, K112I showed half activity compared with the positive control while double mutants of K112I/K172V showed no activity (Fig. 4B).

Here the result indicated that this pair of lysine of *Sc*Alg1-TM work together as a mutual backup and the non-conversed lysine can increase relative activity when the K112I mutant happens, in which case K112I may come to be a lethal mutant. The activity of the mutants showed that Q116L do not have a great effect on the function of *Sc*Alg1-TM (Fig. 4B), which was consistent with the statistics of the relatively long distance and few interactions between Q116 and substrate (Fig. S7).

Considering the anomeric carbon of the terminal sugar residues, the molecular dynamics simulation results showed that Q136 was the closest polar residue to the 4-hydroxyl group of terminal GlcNAc and there were no typical catalytic amino acid residues [44] around it except D48 (Fig. 4B).

### 3.5 Catalytic site and mechanism of *Sc*Alg1

The catalytic site and mechanism were speculated from two aspects: catalytic process and complex structure. Glycosyltransferases catalyze their reactions with either inversion or retention of stereochemistry at the anomeric carbon atom of the donor substrate [45]. Alg1 is an inverting GT in the GT33 family with a characteristic GT-B fold. It consists of two domains with Rossman-like folds connected by a linker. It transfers mannose from GDP-Man to generate a beta-1,4-linked mannose-containing product. Inversion of stereochemistry follows from the fact that the enzyme uses an S_N_2 (substitution nucleophilic bimolecular) reaction mechanism [46] in which an acceptor hydroxyl group attacks the anomeric carbon atom of GDP-Man from one side and UDP leaves from the other.

Typically, enzymes of this type possess an aspartate, glutamate, or histidine residue whose side chain serves to partially deprotonate the incoming acceptor hydroxyl group, rendering it a better nucleophile [47]. However, in the simulation of the complex, it was noticed that the catalytic pocket lacked a basic residue that could act as a catalytic base able to deprotonate the nucleophile residue of the acceptor during the expected S_N_2 chemical reaction. During the simulation, the acetamide part in the side chain of the Q136 residue was near the 4th anomeric carbon of the GlcNAc residue in the terminal of the substrate. Furthermore, Q136 came to be the only polar amino acid except D48 near the GlcNAc part in the pocket, while D48A showed similar activity with wild type (Fig. 4B).

Some structures for GT-B fold inverting glycosyltransferases have been identified in which such a base also appears to be lacking. In these cases, a water-mediated proton-shuttle mechanism has been proposed to achieve the required acceptor deprotonation [48]. It was reported that asparagine could act as the catalytic site for POFUT1, which belongs to GT family 65 and exhibits a GT-B type fold [49].

In order to explore the function of Q136, we mutated it to the amino acid with a similar structure of glutamine: glutamic acid, asparagine and leucine. However, after the activity test in *vitro*, these mutants all showed no activity, which means Q136 is irreplaceable (Fig. 4B).

Summarily, Q136 was predicted to be the catalytic site and the inverting S_N_2 mechanism was also speculated (Fig. 4C), based on MD and reaction mechanism.

### 3.6 The hydrophobic region responsible for the substrate recognition of *Hs*Alg1

*Hs*Alg1 is a membrane protein that is located on the ER membrane [13]. It has no activity after purification and can hardly add sugar residues on the Dol-PP-GlcNAc2 analogue with a short dolichol chain. After analyzing the *Sc*Alg1 and its substrate recognition mechanism, a similar mechanism can be applied to the *Hs*Alg1.

We predicted the structure of the *Hs*Alg1 with AlphaFold2 [50] and then used molecular docking [51,52] to locate the substrate. The N-terminal of the model was different from that of *Sc*Alg1. There are two conformations of GT-B which are opened and closed [53,54]. According to the predicted local distance difference test (pLDDT) of the model, the region of the N-terminal was not accurately predicted, especially the 4^th^ to 6^th^ α helixes part. This region was comprised of a triad of helixes arranged in a triangular configuration. Within this structure, a substrate recognition channel was present in *Sc*Alg1. Notably, this channel was absent in the predicted structure of *Hs*Alg1 by AlphaFold2 (Fig. 5A, S8).

**Figure 5.**
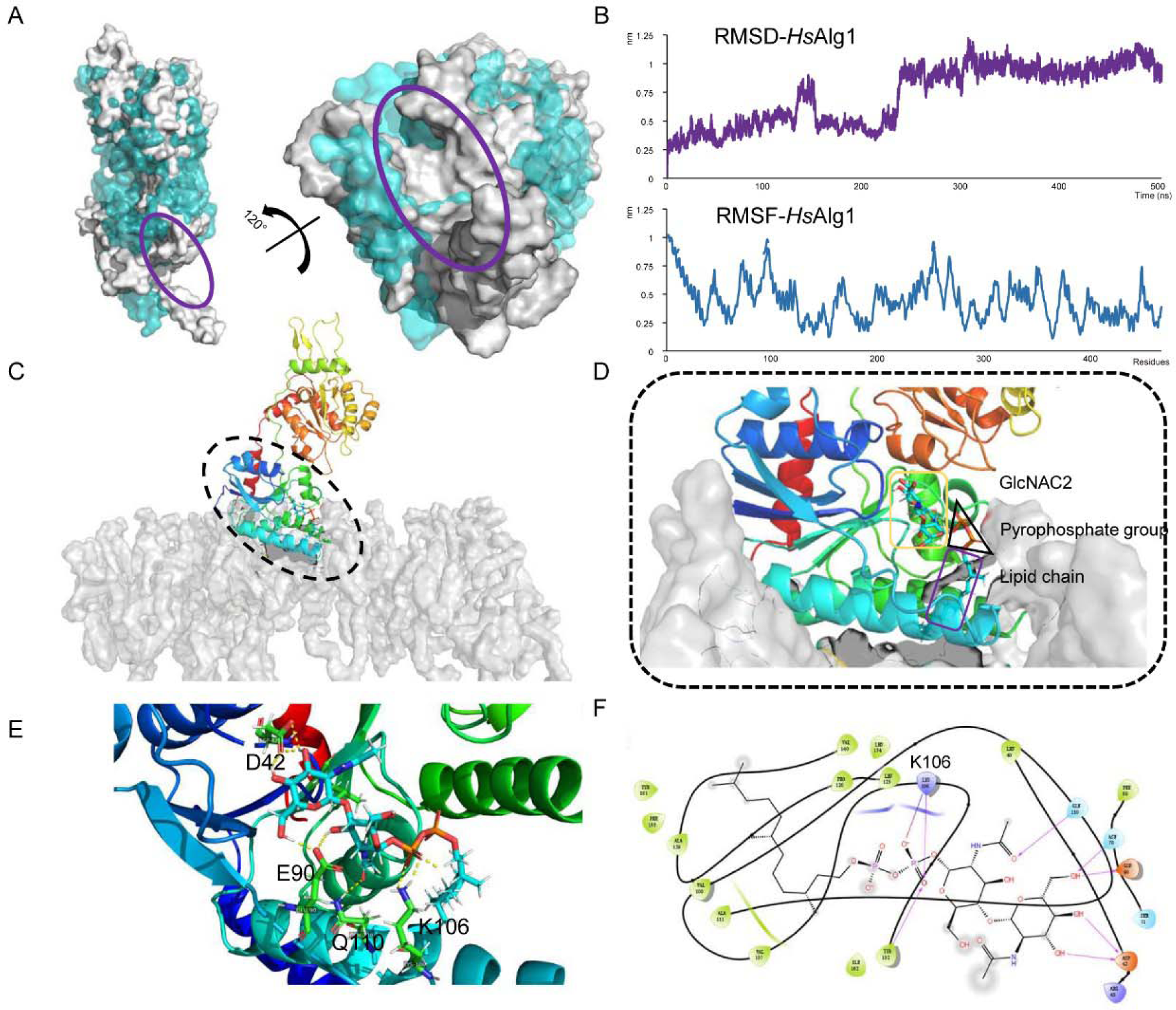
The substrate recognition region of *Hs*Alg1. (A) The structure of *Hs*Alg1 before (blue) and after (grey) MD. The hydrophobic groove appeared after MD. (B) The RMSD and RMSF of the MD. (C) *Hs*Alg1 was located on the membrane during MD. (D) The substrate was fixed in the interface between *Hs*Alg1 and the membrane. (E) The interactions of *Hs*Alg1 and Dol2-PP-GlcNAc2. (F) The salt bridge formed between K106 and PPGn2.

To improve the structure of *Hs*Alg1, we added the model of *Hs*Alg1 on the membrane [31] and the surrounding solvent to simulate the environment in *vivo*. After building the system, molecular dynamics simulation was applied to reduce the inaccuracy. Due to the specificity of structural changes and membrane protein, 500 ns simulation was applied and the change of RMSD was observed (Fig. 5B). As expected, the channel opened after simulation and the substrate binding area was exposed (Fig. 5A, S8).

Apart from forming the hydrophobic channel, three helixes were anchored to the membrane. Together with the first transmembrane domain, these three helixes located *Hs*Alg1 on the ER membrane (Fig. 5C).

With the relatively open protein conformation, the complex of *Hs*Alg1, Dol2-PP-GlcNAc2 and GDP-Man was assembled, with the bilayer membrane built by CHARMM-GUI [32,55]. The model was inserted into the membrane with the appropriate angle predicted by the OPM server [31]. After 100 ns simulation, the dolichol chain was in the channel and the sugar part was in the pocket (Fig. 5D). The pyrophosphate part of the substrate formed salt bridges with the K106. This lysine was conserved and was the homologous amino acid of K112 in *Sc*Alg1 (Fig. 3F), which was proved to be the key site for substrate recognition. In this case, the hydrophobic groove was proved to be the region responsible for substrate recognition again.

When the final 1 ns was picked during the simulation of the complex, the location of the substrate was focused. The dolichol chain was in the channel which was surrounded by the three helixes and lipids of the membrane (Fig. 5D). The pyrophosphate part was in the interface of the channel and the membrane, which was partially mixed with phosphoric acid parts of the phospholipids (Fig. 5E). The sugar terminal was stretched into the pocket of the *Hs*Alg1.

This hydrophobic region served two biological purposes. Most of the region was buried in the membrane, which both located the *Hs*Alg1 and attracted the dolichol chain of the substrate. Docking with PPGn2 to compare with the substrate recognition of *Sc*Alg1-TM, there were basic amino acids that interacted with the pyrophosphate portion of the substrate to fix the substrate at the top of this region in *Hs*Alg1 (Table S3, Fig. 5F). By maintaining a suitable distance from the surface of the phospholipid bilayer, lysine was able to pick up the substrates from a large number of phospholipid molecules.

In the exploration of the mechanism of substrate recognition, the hydrophobic channel attracted the substrate by the hydrophobic interaction with the dolichol and then the lysines fixed the substrate with the salt bridge. This region anchored the protein to the membrane and identified substrates floating within the endoplasmic reticulum. This synergistic effect was found in both *Sc*Alg1 and *Hs*Alg1, even in the other GT-B of the N-glycosylation. During the evolution of these GTs with a substrate of similar structure, this hydrophobic region with the basic amino acid on the side might be conserved.

## 4. Conclusion

This study reported the molecular mechanism of substrate recognition and the catalytic mechanism of Alg1. In the activity tests in *vitro*, it was found that *Sc*Alg1 can use GDP-Alt as its sugar donor with a similar preference to its natural donor GDP-Man. By molecular docking and simulation, a series of residues in the C-terminal Rossmann fold were found to help recognize sugar donors, which revealed the molecular mechanism of non-natural donor recognition. The hydrophobic channel recruited the dolichol of the substrate to the hydrophobic groove, with the synergistic effect of two lysines forming salt bridges with the pyrophosphate group to fix the substrate. The reaction of *Sc*Alg1 follows an S_N_2 mechanism, as in classical inverting glycosyltransferases, but a glutamine residue rather than a basic residue deprotonates the hydroxyl group of the nucleophile. The hydrophobic channel of *Hs*Alg1 opened after the simulation of the system and the region for substrate recognition was speculated. Also, the dolichol part of the substrate was proven to promote substrate binding. Examining the enzymatic characteristics of *Hs*Alg1, encompassing substrate recognition mechanisms and preferences bears substantial implications for comprehending Alg1-CDG. This endeavour facilitates the elucidation of disease pathogenesis, pinpointing prospective therapeutic targets, devising personalized treatment modalities, and advancing pharmaceutical development, thereby potentially affording enhanced interventions for this rare disorder.

## Declaration of competing interest

The authors declare that they have no known competing financial interests or personal relationships that could have appeared to influence the work reported in this paper.

## Declaration of competing interest

The authors have no competing interests.

## Acknowledgements

This study was financially supported by the National Natural Science Foundation of China

## Appendix A. Supplementary data

Movie 1.

Movie 2.

Download: Download Word document

Supplementary material

## Data availability

Data will be made available on request.

## Notes

### Competing Interest Statement

The authors have declared no competing interest.

